# Evolution of asymmetric gamete signaling and suppressed recombination at the mating type locus

**DOI:** 10.1101/518654

**Authors:** Zena Hadjivasiliou, Andrew Pomiankowski

## Abstract

The two partners required for sexual reproduction are rarely the same. This pattern extends to species which lack sexual dimorphism yet possess self-incompatible gametes determined at mating-type regions of suppressed recombination, likely precursors of sex chromosomes. Here we investigate the role of cellular signaling in the evolution of mating-types. We develop a model of ligand-receptor dynamics within cells, and identify factors that determine the capacity of cells to send and receive signals. The model specifies conditions favoring the evolution of gametes producing ligand and receptor asymmetrically and shows how these are affected by recombination. When the recombination rate can evolve, the conditions favoring asymmetric signaling also favor tight linkage of ligand and receptor loci in distinct linkage groups. These results suggest that selection for asymmetric signaling between gametes was the first step in the evolution of non-recombinant mating-type loci, paving the road for the evolution of anisogamy and sexes.

## 1 Introduction

Sex requires the fusion of two cells. With few exceptions, the sexual process is asymmetric with partnering cells exhibiting genetic, physiological or behavioral differences. The origins of sexual asymmetry in eukaryotes trace back to unicellular organisms with isogametes lacking any size or mobility difference in the fusing cells [1, 2, 3, 4, 5, 6]. Isogamous organisms are divided into genetically distinct mating types, determined by several mating type specific genes that reside in regions of suppressed recombination [7, 8, 9, 10]. The morphologically identical gametes mate disassortatively, scarcely ever with members of the same mating type. It follows that only individuals of a different mating type are eligible mating partners. This arrangement poses a paradox as it restricts the pool of potential partners to those of a different mating type, introducing a major cost [4].

Several hypotheses have been proposed to explain the evolution of isogamous mating types [11, 12, 13]. Mating types could serve as a restrictive mechanism preventing matings between related individuals thereby avoiding the deleterious consequences of inbreeding [14, 15, 16]. Another idea is that mating types facilitate uniparental inheritance of mitochondria, which leads to improved contribution of the mitochondrial genome to cell fitness [17, 18, 19, 20, 21, 22, 23, 24]. Both hypotheses have been studied extensively and offer compelling arguments. Nevertheless, the existence of several species where inbreeding [13, 12] or biperental inheritance of mitochondria [25, 12] are the rule but nonetheless maintain mating types, indicates that these ideas may not alone explain the evolution of mating types.

An alternative hypothesis is that mating types are determined by the molecular system regulating gamete interactions [26, 27, 4]. Such interactions dictate the success of mating by guiding partner attraction and recognition and the process of cell fusion, and have been shown to be more efficient when operating in an asymmetric manner [26]. For example, diffusible molecules are often employed as signals that guide synchronous entry to gametogenesis or as chemoatractants [28, 29, 30, 31]. Secreting and sensing the same diffusible molecule impedes the ability of cells to accurately detect external signals and makes partner finding many-fold slower [26]. In addition, secreting and detecting the same molecule in cell colonies can prevent individuals responding to signals from others [32]. Our previous review revealed that sexual signaling and communication in isogamous species are universally asymmetric [27]. This applies throughout the sexual process from signals that lead to gametic differentiation, to attraction via diffusible pheromones and interactions via surface bound molecules during cell fusion [27].

In this work we take this analysis further by explicitly considering ligand-receptor interactions between and within cells. We directly follow the dynamics of ligand and receptor molecules that are surface bound and determine the conditions under which the production of within cell ligand-receptor pairs impedes between cell communication. We use this framework to explore the evolution of gametic interactions and show that asymmetric signaling roles and tight linkage between receptor and ligand loci both evolve due to selection for robust intercellular communication and quick mating. Our findings demonstrate that the evolution of mating type loci with suppressed recombination can be traced back to the fundamental selection for asymmetric signaling during sex.

## 2 Theoretical set-up

Consider a population where cells encounter one another at random and can mate when in physical contact. Interactions between cells leading to successful mating are dictated by a ligand-receptor pair. Population wide effects may emerge if the ligand is highly diffusible [26, 32]. The employment of membrane bound ligands during sexual signaling is universal, whereas diffusible signals are not [27]. In this work we therefore assume that the ligand-receptor interactions only operate locally. Receptors remain bound to the cell surface and ligands only undergo localized diffusion (Fig. 1) as is the case in several yeast and other unicellular eukaryotes [33, 29, 34, 35]. The following equations describe the concentration of free ligand *L*, free receptor *R* and bound ligand *LR* within a single cell,

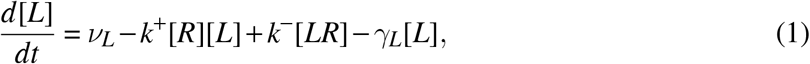

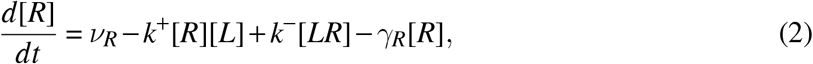

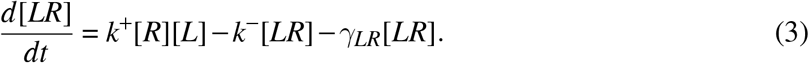

*ν_L_* and *ν_R_* describe the rate of production of the ligand and receptor respectively. *γ_L_, γ_R_*, and *γ_LR_*, are the degradation rate of the ligand, receptor and bound complex respectively. The terms *k*^+^ and *k*^−^ are the binding and unbinding rates that determine the affinity of the ligand to its receptor within a single cell. We can solve Eq. (1–3) by setting the dynamics to zero to obtain the amount of free ligand, free receptor ([*L*]*, [*R*]*) and bound complex at steady state ([*LR*]*),

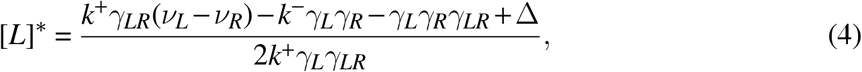

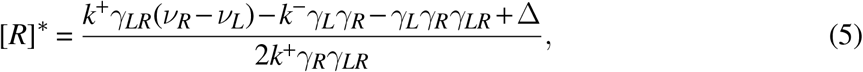

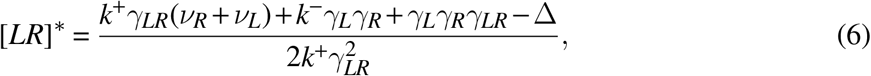

**Figure 1:**
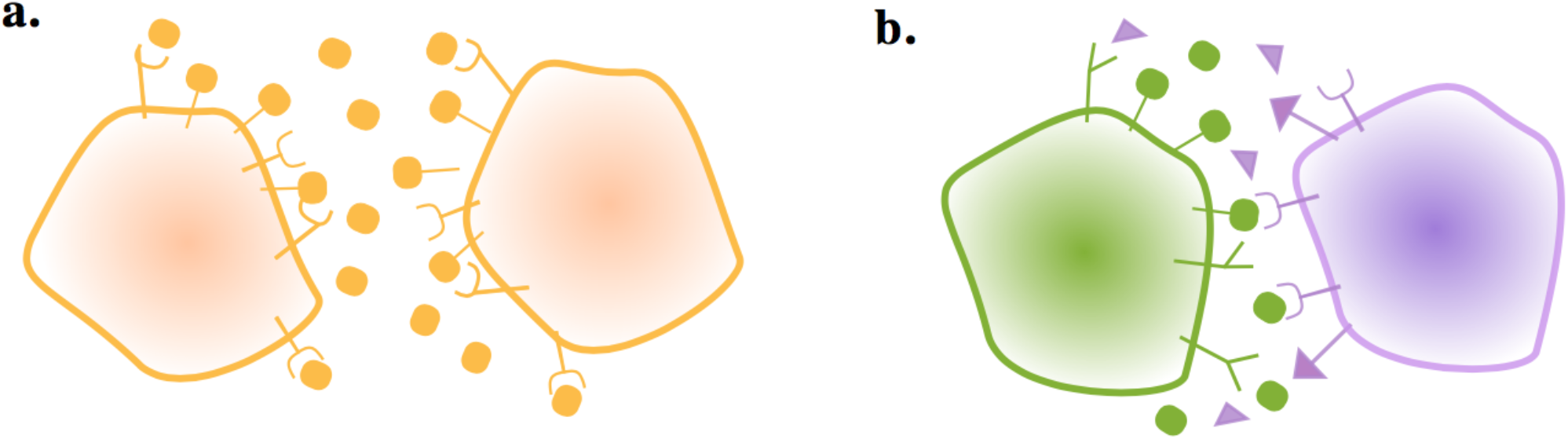
Gametes communicate through ligand and receptor molecules. The ligand can be either membrane bound or released in the local environment. (a) When the interacting cells produce ligand and receptor symmetrically, the ligand will bind to receptors on its own membrane as well as those on the other cell. This may impair intercellular signaling. (b) Producing the ligand and receptor in an asymmetric manner resolves this issue.

Where Δ is given by,

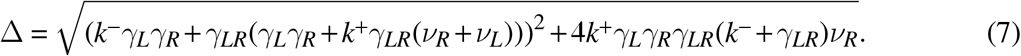

We assume that the timescale of encounters and interactions between cells is longer than the timescale of ligand and receptor production and degradation. Hence the concentrations of [*L*], [*R*] and [*LR*] in individual cells will be at steady state when two cells meet. The likelihood of a successful mating between two cells depends not just on partner signaling levels but also on how accurately the cells can compute the signal produced by their partner. Binding of ligand and receptor originating from the same cell can obstruct this interaction. To capture this, we define the strength of the incoming signal for cell_1_ when it interacts with cell_2_ as,

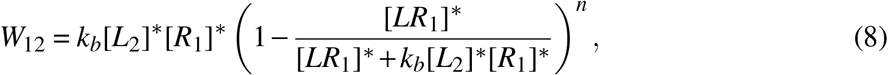

where subscripts denote concentrations in cell_1_ and cell_2_, and the parameter *k_b_* determines the affinity of the ligand and receptor between cells. If *k_b_* is the same as the affinity of receptor and ligand within cells, then 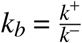. We also consider cases where 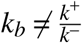, for example, when ligand interacts differently with receptors on the same as opposed to a different cell [36, 37].

The cost of self-signaling is determined by *n*. When *n* = 0, *W*_12_ reduces to *k_b_*[*R*_1_]*[*L*_2_]* with the incoming signal dependent on the concentration of ligand produced by cell_2_ and receptor produced by cell_1_. This corresponds to a case where self-binding does not lead to activation but only causes an indirect cost through the depletion of available ligand and receptor molecules. When *n* ≥ 1, binding within a cell leads to some form of activation that interferes with between cell signaling, imposing a cost in evaluating the incoming signal. Higher values for *n* correspond to more severe costs due to self-binding.

The likelihood that two cells successfully mate (*P*) depends on the quality of their interaction given by,

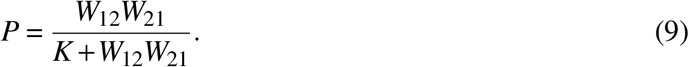

Eq. (9) transforms the signaling interaction into a mating probability. For the analysis that follows, we choose large values of *K* so that *P* is far from saturation and depends almost linearly on the product *W*_12_*W*_21_. In summary, the probability that two cells mate is defined by the production and degradation rates of the ligand and receptor molecule, and the binding affinities between and within cells.

### 2.1 Evolutionary model

To explore the evolution of signaling roles, we simplify the model by assuming that the degradation rates *γ_L_,γ_R_,γ_LR_* are constant and equal to *γ*, and investigate mutations that quantitatively modify the ligand and receptor production rates. Consider a finite population of *N* haploid cells where loci that control ligand and receptor production are quantitative traits. The ligand and receptor production rates of cell_*i*_ is denoted by (*ν_L_i__, ν_R_i__*). We also consider different versions of the ligand and its receptor. Cells have two ligand-receptor pairs, (*L,R*) and (*l, r*) which are mutually incompatible, so the binding affinity is zero between *l* and *R*, and between *L* and *r*. Each cell has a (*L, R*) and (*l, r*) state, which are subject to mutational and evolutionary pressure as described below. *W*_12_ is re-defined as the summation of the interactions of these two ligand-receptor pairs,

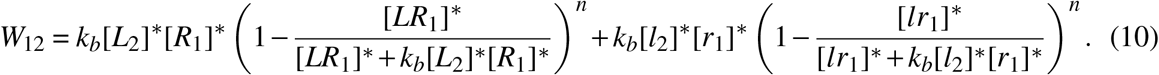

Again for the sake of simplicity, the ligand-receptor affinities are set to be the same between and within cells for each ligand-receptor pair (i.e. *k*^+^, *k*^−^ and *k_b_* are the same for *L − R* and *l − r* interactions). A cell undergoes mutation that changes 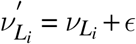 with *ϵ* ~ *N*(0, *σ*) with probability *μ*. The same mutational process occurs for all ligand and receptor production rates. We assume that mutation occurs independently at different loci and that there is a maximum capacity for ligand and receptor production, so that *ν_L_ + ν_l_* < 1 and *ν_R_ + ν_r_* < 1. It follows that the production rates in the two ligand genes are not independent of one another and similarly for the two receptor genes.

We also consider cases where *ν_L_ + ν_l_* < *α* and *ν_R_ + ν_r_* < *α* for *α* ≠ 1 to reflect synergy (*α* > 1) or competition (*α* < 1) between the production of the two ligands (or receptors). For example, synergy between two ligands (or receptors) could reflect reduced energy expenditure for the cell if the same machinery is used to produce the two molecules. Competition on the other hand could reflect additional costs due to the production of two different ligands (or receptors).

Selection on ligand-receptor production rates is governed by the likelihood that cells pair and produce offspring. We assume that cells enter the sexual phase of their life cycle in synchrony, as is the case in the majority of unicellular eukaryotes [27]. Pairs of cells are randomly sampled (to reflect random encounters) and mate with probability *P* defined in Eq. (9). Cells failing to mate are returned to the pool of unmated individuals. The process is repeated until *M* cells have mated. Each mated pair produces 2 haploid offspring so the population size shrinks from *N* to *M*. The population size is restored back to *N* by sampling with replacement. It follows that Eq. (9) and together provide a proxy for fitness according to the ligand and receptor production rates of individual cells. Initially, recombination is not allowed between the genes controlling ligand and receptor production but then is considered in a later section.

## 3 Results

### 3.1 Dependence of gamete interactions on physical parameters

The strength of an incoming signal *W*_12_ depends on the concentration of free receptor in cell_1_ and free ligand in cell_2_, and the cost of self-binding (*n*) (Eq. (10)). The steady state concentration of [*L*], [*R*] and [*LR*] are governed by different production rates (Fig. S1; details of the derivation can be found in the Methods section). For low degradation rates (*γ* small), the removal of available molecules is dominated by self-binding (*k*^+^) (Eq. (1) and (2) and Fig. 2a, b). At the same time, a lower degradation rate leads to higher levels of ligand and receptor (Fig. 2a) even if the relative drop of free ligand and receptor is steeper as *k*^+^ increases (Fig. 2b). As a consequence, the ability of a cell to generate a strong signal and read incoming signals can change drastically when the pair of interacting cells produce the ligand and receptor in a symmetric manner (e.g. (*ν_L_, ν_R_, ν_l_, ν_r_*) = (1,1,0,0) for both cells) rather than in an asymmetric manner (e.g. (*ν*_*L*_1__, *ν*_*R*_1__, *ν*_*l*_1__, *ν*_*r*_1__) = (1,0,0,1) and (*ν*_*L*_2__, *ν*_*R*_2__, *ν*_*l*_2__, *ν*_*r*_2__) = (0,1, 1, 0)). The fold-increase in *W*_12_ is large even when self-binding confers no cost (*n* = 0), while larger values for *n* ramp up the costs (Fig. 2c). If cells produce the ligand and receptor asymmetrically, self-binding ceases to be a problem in receiving incoming signals.

**Figure 2:**
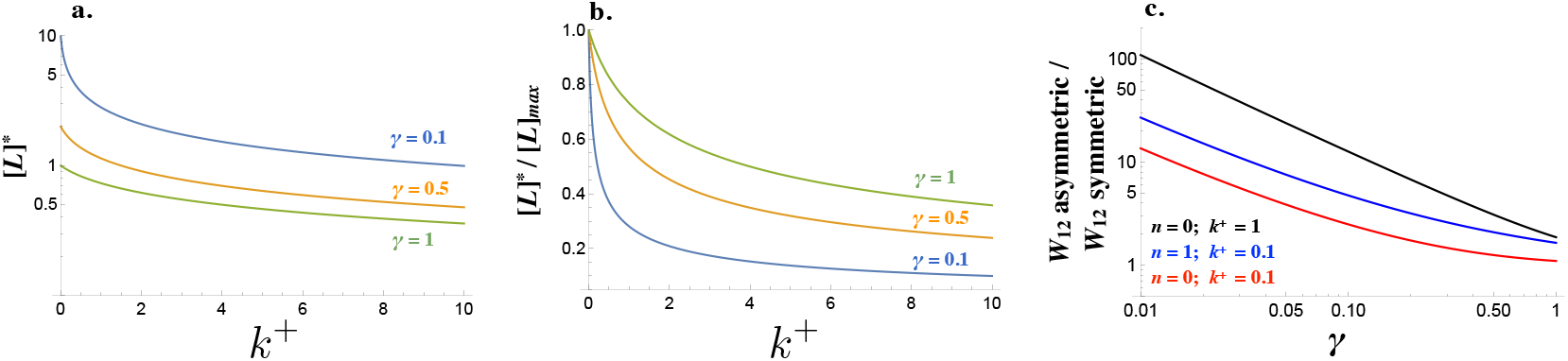
Signaling interactions between mating cells can be severely impaired due to ligand-receptor interactions in the same cell. (a) The amount of free ligand in individual cells at steady state [*L*]* and (b) normalized amount of free ligand at steady state [*L*]*/[*L*]_*max*_ varies with the intracellular binding rate *k*^+^ and degradation rate *γ*. (c) The relative amount of incoming signal *W*_12_ for a cell that produces ligand and receptor asymmetrically versus symmetrically decreases with the degradation rate *γ* and weaker binding *k*^+^. Other parameters used: *n* =1, *k*^−^ = 1, *k_b_* = 1.

Although the strength of the signaling interaction between two cells (*W*_12_*W*_21_) may improve when the interacting cells produce the ligand and receptor asymmetrically, this need not be the case.

Consider the interaction of a resident cell with production rates (*ν_L_, ν_R_, ν_l_, ν_r_*)_*res*_ = (1,1,0,0) with itself and a mutant cell with production rates given by (*ν_L_, ν_R_, ν_l_, ν_r_*)_*mut*_ = (1 − *dx*, 1 − *dy, dx, dy*). For all values of *dx* and *dy*, [*W*_12_*W*_21_]_*res+mut*_ − [*W*_12_*W*_21_]_*res+res*_ < 0 (Fig. 3a). It follows that (*ν_L_, ν_R_, ν_l_, ν_r_*) = (1, 1, 0, 0) cannot be invaded by any single mutant.

**Figure 3:**
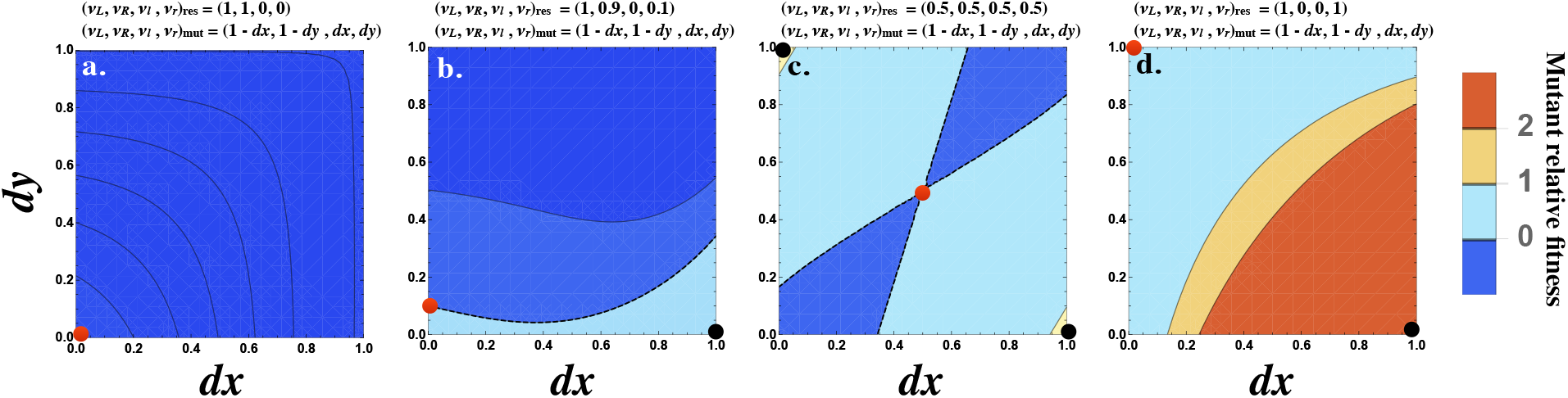
Fitness advantage of rare mutations conferring signaling asymmetry. The fitness of a rare mutant is plotted relative to the resident [*W*_12_*W*_21_]_*res+mut*_ − [*W*_12_*W*_21_]_*res+res*_. The production rate of the mutant cell is (*ν_L_, ν_R_, ν_l_, ν_r_*)_*mut*_ = (1 − *dx*, 1 − *dy, dx, dy*), where *dx* and *dy* are plotted on the *x* and *y* axes respectively. The resident production rate (*ν_L_, ν_R_, ν_l_, ν_r_*)_*res*_ is shown as a red dot and varies (a) (1,1,0,0)_*res*_, (b) (1,0.9,0,0.1)_*res*_, (c) (0.5,0.5,0.5,0.5)_*res*_ and (d) (1,0,0,1)_*res*_. The mutant (*dx, dy*) with maximum fitness is shown as a black dot. The contour where [*W*_12_*W*_21_]_*res+mut*_ = [*W*_12_*W*_21_]_*res+res*_ is marked by a black dashed line (b and c). The fitness difference is always negative in (a) and always positive in (d). Other parameters used: *n* =1,*γ* = 0.5, *k*^+^ = 1, *k*^−^ = 1, *k_b_* = 1.

However, if the resident is already slightly asymmetric so that (*ν_L_, ν_R_, ν_l_, ν_r_*)_*res*_ = (1,0.9,0,0.1) then a mutant conferring an asymmetry in the opposite direction can be better at interacting with the resident (Fig. 3b). When the resident produces both ligand and receptor equally (i.e. (*ν_L_, ν_R_, ν_l_, ν_r_*)_*res*_ = (0.5,0.5,0.5,0.5)) then all mutants are favored, with the strongest interaction occurring with mutants that produce the ligand or receptor asymmetrically (i.e. (*ν_L_, ν_R_, ν_l_, ν_r_*)_*mut*_ = (1,0,0,1) or (0, 1, 1, 0) are (Fig. 3c). Finally, when the resident production rates are already strongly asymmetric given by (*ν_L_, ν_R_, ν_l_, ν_r_*)_*res*_ = (1,0,0,1), a mutant with an asymmetry in the opposite direction is most strongly favored (Fig. 3d). Note that a population composed only of cells with production rates at (*ν_L_, ν_R_, ν_l_, ν_r_*)_*res*_ = (1,0,0,1) is not viable since the probability that two such cells mate is zero. However, this analysis provides insight about how asymmetry in signaling evolves.

### 3.2 Evolution of mating types with asymmetric signaling roles

To explore the evolution of signaling asymmetry, we follow mutations that alter the relative production of two mutually incompatible types of ligand and receptor (*L,R*) and (*l, r*). To ease understanding, the population symmetry *s* in the production of ligand and receptor is measured,

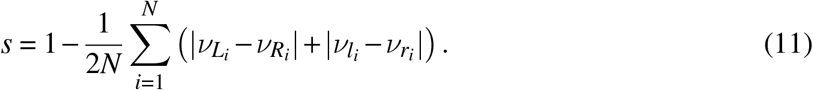

The population is symmetric (*s* = 1) if cells produce ligand and receptor equally, for both types (i.e. (*ν_R_, ν_L_, ν_r_, ν_l_*) = (*a, a*, 1 − *a*, 1 − *a*), for constant *a*), and fully asymmetric (*s* = 0) when cells adopt polarized roles (i.e. (*ν_L_, ν_R_, ν_l_, ν_r_*) = (1,0,0,1) or (0,1,1,0)).

Starting from a population where all cells are symmetric producers of only one ligand and receptor, (*ν_L_, ν_R_, ν_l_, ν_r_*) = (1,1,0,0), the population evolves to one of two equilibria (Fig. 4a). *E*_1_ where *s*\* ≈ 1 and all cells produce the ligand and receptor symmetrically (*ν_L_, ν_R_, ν_l_, ν_r_*) ≈ (1,1,0,0) or *E*_2_ where *s*\* ≈ 0 and the population is divided into ligand and receptor producing cells, with equal frequencies of (*ν_L_, ν_R_, ν_l_, ν_r_*) ≈ (1,0,0,1) and (*ν_L_, ν_R_, ν_l_, ν_r_*) ≈ (0,1,1,0) (Fig. 4b, c). Equilibria with intermediate values of *s*\* are not found. The exact production rates at *E*_1_ and *E*_2_ exhibit some degree of noise due to mutation and finite population size (Fig. 4b, c). At *E*_2_, individual cells with high *ν_R_* (and low *ν_r_*) have low *ν_L_* (and high *ν_l_*), confirming that *s*\* ≈ 0 captures a fully asymmetric steady state.

**Figure 4:**
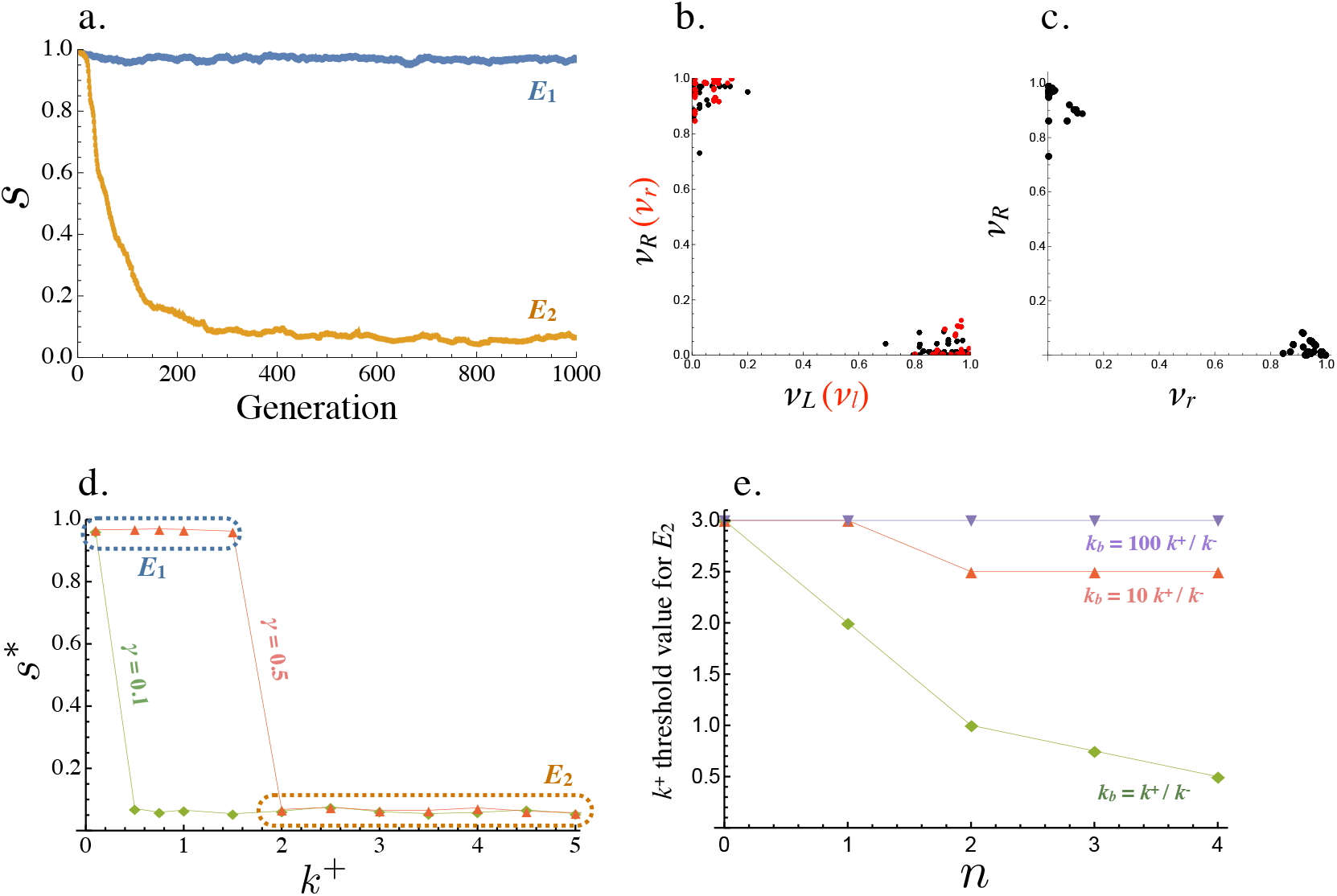
Evolution of asymmetric signaling. (a) An example of evolution to the two signaling equilibria, *E*_1_ (*s* =1 full symmetry when *k*^+^ =1) and *E*_2_ (*s* = 0 full asymmetry when *k*^+^ = 5). (b) Production rates of individual cells in the population for the receptor-ligand pairs *L − R* (black) and *l − r* (red) at *E*_2_. (c) Production rates of individual cells for the two receptor types *R* and *r* at *E*_2_. (d) Steady state signaling symmetry *s*\* against the intracellular binding rate (*k*^+^) for different degradation rates (*γ*). (e) Threshold value of *k*^+^, beyond which *E*_2_ evolves from *E*_1_, plotted versus the cost of self-binding (*n*). The relationship is shown for different values of strength of between cell signaling (*k_b_*) relative to strength of within cell signaling (*k*^+^/*k*^−^). Other parameters used in numerical simulations are given in the Supplemental Material.

Whether *E*_2_ is reached from *E*_1_ depends on key parameters that determine the strength of self-binding and signaling interactions between cells. *E*_1_ persists and no asymmetry evolves when *k*^+^ (the intracellular ligand-receptor binding coefficient) is small (Fig. 4d). In this case, the concentration of self-bound ligand-receptor complex is small (Eq. (6)) and there is little cost of self-signaling (Eq. (8)), so there is weak selection in favor of asymmetry. The opposite is true for larger values of *k*^+^, as self-binding now dominates and restricts between cell signaling, promoting the evolution of asymmetry (Fig. 4d). The transition from *E*_1_ to *E*_2_ occurs at a smaller value of *k*^+^ when the degradation rate (*γ*) is decreased (Fig. 4d), as the effective removal of free ligand and receptor depends more strongly on intercellular binding (Fig. 2a, b).

Another important consideration is the relative strength of signaling within and between cells, given by *k*^+^/*k*^−^ and *k_b_* respectively. For example, the threshold value of the within cell binding rate beyond which symmetric signaling (*E*_1_) evolves to asymmetric signaling (*E*_2_, Fig. 4a) increases when *k_b_* becomes much larger than *k*^+^/*k*^−^ (Fig. 4e). Furthermore, this threshold value is smaller for larger values of *n* indicating that asymmetric signaling is more likely to evolve when the cost for self-signaling is higher (larger *n*, Fig. 4e). However, asymmetric signaling can evolve even when self-binding carries no cost (*n* = 0) as high rates of self-binding can restrict the number of ligand and receptor molecules free for between cell interactions (Fig. 4e).

These observations suggest that both *E*_1_ and *E*_2_ are evolutionary stable states and the transition from *E*_1_ to *E*_2_ depends on the mutational process, drift and the parameters that determine signaling interactions. To explore this we investigated the stability of *E*_1_ in response to rare mutations in the receptor and ligand production rates. We assume the population is initially at *E*_1_ (i.e. (*ν_L_, ν_R_, ν_l_, ν_r_*) = (1,1,0,0)), introduce mutations in the receptor and ligand loci (*ν_L_, ν_R_, ν_l_, ν_r_*) = (1 − *dx*, 1, *dx*, 0) and (*ν_L_, ν_R_, ν_l_, ν_r_*) = (1,1 − *dy*, 0, *dy*) at frequency *μ_a_*, and calculate the population symmetry at steady state for different values of *dx* and *dy* (Fig. 5). Single mutations never spread (i.e. if *dx* = 0 no value of *dy* allows mutants to spread and vice versa). This is in agreement with the analytical predictions presented in the previous section (Fig. 3). When both *dx* and *dy* are nonzero the population may evolve to *E*_2_, where the two mutants reach equal frequencies at ~0.5 and replace the resident. The basin of attraction for *E*_2_ (and so asymmetric signaling roles) is larger when *k*^+^ and *μ_a_* are high and *γ* is small (Fig. 5a-d), as predicted analytically (Fig. 2, 3) and in accordance with our findings when mutations were continuous (Fig. 4). Note that the magnitude of the mutation rates matters in our system. Single mutations can be slightly deleterious (as predicted analytically, Fig. 3a), but the presence of mutants that are asymmetric in opposite directions gives rise to positive epistatic effects (Fig. 3b-d). This explains why smaller values of *μ_a_* result in narrower basins of attraction for *E*_2_ (Fig. 5).

**Figure 5:**
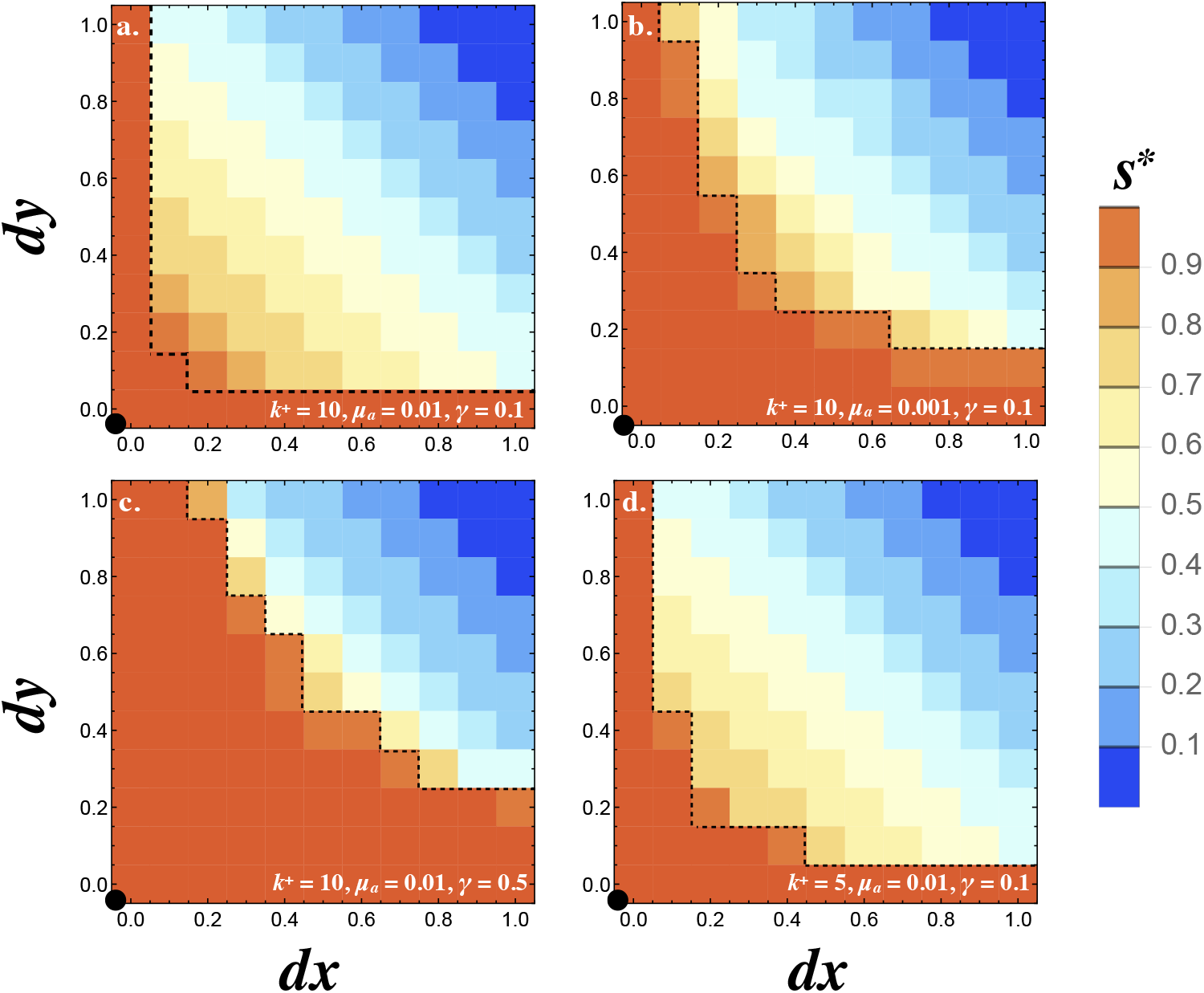
Invasion of *E*_1_. Contour plots showing the steady state degree of symmetry (*s*\*) in a population with resident (*ν_R_, ν_L_, ν_r_, ν_l_*) = (1,1,0,0). Two mutations are introduced (1 − *dx*, 1, *dx*, 0) and (1,1 − *dy*, 0, *dy*) at rate *μ_a_* and their fate is followed until they reach a stable frequency. Orange contours outside the dotted line show the region where both mutants are eliminated and the resident persists (*s*\* = 1). All other colors indicate that the two mutants spread to equal frequency 0.5 displacing the resident (*s*\* < 1). The degree of signaling symmetry at equilibrium is dictated by the magnitude of the mutations given by *dx* and *dy*. The different panels show (a) between cell signaling *k*^+^ = 10, mutation rate *μ_a_* = 0.01 and degradation rate *γ* = 0.1, (b) higher mutation rate *μ_a_* = 0.001, (c) high degradation rate *γ* = 0.5 and (d) weaker between cell signaling *k*^+^ = 5. The resident type is marked by a black dot at the origin. The dashed line marks the regions above which the two mutants spread to displace the resident and reach a polymorphic equilibrium at equal frequencies. Other parameters used and simulation details are given in the Supplementary Material.

We next investigated how mutations invade when the resident already signals asymmetrically (i.e. produces both ligands). The resident was set to (*ν_L_, ν_R_, ν_l_, ν_r_*)_*res*_ = (1 − *dx*, 1, *dx*, 0) and a mutant able to produce both receptors (*ν_L_, ν_R_, ν_l_, ν_r_*)_*mut*_ = (1,1 − *dy*, 0, *dy*) was introduced. If *dx* = 0, a mutant conveying a small asymmetry in receptor production increases in frequency until the population reaches a polymorphic state with the resident and mutant at 50% (Fig. 6a). If on the other hand the resident exhibits an asymmetry but the mutant does not (i.e. *dy* = 0 and *dx* > 0), the mutant replaces the resident. It follows that an asymmetry in both ligand and receptor production is necessary for the evolution of a signaling asymmetry as predicted analytically (Fig. 3). We also consider a resident type that produces both ligands and both receptors with some degree of asymmetry in ligand production (i.e. (*ν_L_, ν_R_, ν_l_, ν_r_*)_*res*_ = (0.5 − *dx*, 0.5,0.5 + *dx*, 0.5)) and map the spread of a mutant with asymmetry is receptor production (*ν_L_, ν_R_, ν_l_, ν_r_*)_*mut*_ = (0.5,0.5 − *dy*, 0.5,0.5 + *dy*). The pairwise invasability plots for values of *dx* and *dy* show that signaling asymmetries in opposite directions are favored. These evolve to a polymorphic state with equal frequencies of cells at *dx* = *dy* = −0.5 and *dx* = *dy* = 0.5 (Fig. 6b). These findings together illustrate how the asymmetric state *E*_2_ evolves from the symmetric state *E*_1_.

**Figure 6:**
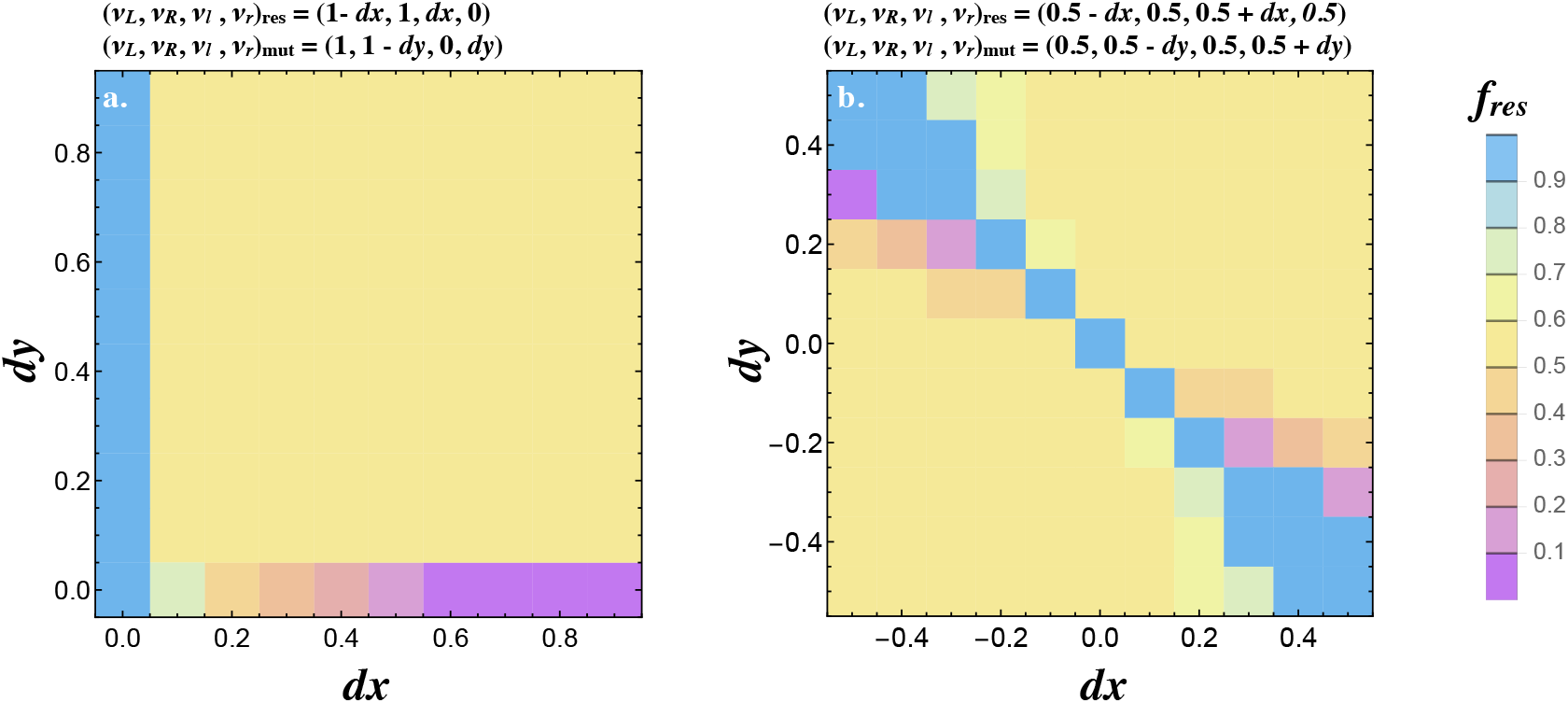
Joint evolution of receptor and ligand asymmetry. Contour plots show the equilibrium frequency of a resident with production rates (*ν_L_, ν_R_, ν_l_, ν_r_*)_*res*_ = (1 − *dx*, 1, *dx*, 0) (a) (*ν_L_, ν_R_, ν_l_, ν_r_*)_*res*_ = (0.5 − *dx*, 0.5,0.5 + *dx*, 0.5) (b), following a mutation (*ν_L_, ν_R_, ν_l_, ν_r_*)_*mut*_ = (1,1 − *dy*, 0, *dy*) (a) and (*ν_L_, ν_R_, ν_l_, ν_r_*)_mut_ = (0.5,0.5 − *dy*, 0.5,0.5 + *dy*) (b). The mutant is introduced at a frequency *μ_a_* = 0.01. Other parameters used and simulations details are given in the Supplemental Material.

Finally, we wondered how synergy or competition between the two ligands (or receptors) could affect our results. When the two ligands (or receptors) exhibit synergy so that *ν_L_* + *ν_l_* < *α* and *ν_R_* + *ν_r_* < *α* for *α* > 1, a signaling asymmetry evolves more easily (for smaller values of *k*^+^, Fig. S3). Now the second ligand (or receptor) begins to evolve without imposing a cost on the preexisting ligand (or receptor) and can therefore remain present in the population longer until an asymmetry in the opposite direction evolves in other cells. The reverse dynamics are observed when the two ligands (or receptors) compete with one another (*ν_L_* + *ν_l_* < *α* and *ν_R_* + *ν_r_* < *α* for *α* < 1) (Fig. S3).

### 3.3 Effects of recombination

The results above assume that the loci controlling ligand and receptor production are tightly linked which prevents the production of deleterious combinations following meiosis. Recombination is a minor problem at the *E*_1_ equilibrium which is monomorphic (except for mutational variation). But it is likely to be a problem at the polymorphic *E*_2_ equilibrium. At *E*_2_, mating between (*ν_L_, ν_R_, ν_l_, ν_r_*) = (1,0,0,1) and (0, 1, 1, 0) cells generates non-asymmetric recombinant ligand-receptor combinations, either (1,1,0,0) or (0,0,1,1). To implement recombination we assume that the two ligands are tightly linked in a single locus and are inherited as a pair (likewise the two receptors), and investigate the effects of recombination between the ligand and receptor loci.

Consider the effect of recombination on a population at *E*_1_. As before, the population either stays at *E*_1_ or evolves to *E*_2_ dependent on parameter values (Fig. 7a). When the population evolves to *E*_2_, *s*\* becomes larger as the recombination rate (*ρ*), increases (Fig. 7 b). For low recombination rates (*ρ* ≤ 0.1), the population largely consists of equal frequencies of (1,0,0,1) and (0, 1, 1, 0) cells, producing the ligand and receptor asymmetrically. A small percentage of recombinant cells produce conspecific pairs of ligand and receptor (*ν_L_, ν_R_, ν_l_, ν_r_*) = (1,1,0,0) and (0,0,1,1) (Fig. 7b, c). Recombination in this case creates “macromutations” where production rates that were 0 become 1 and vice versa. As the recombination rate rises (*ρ*≥0.2), the two leading cell types diverge from (*ν_L_, ν_R_, ν_l_, ν_r_*) = (1,0,0,1) and (0, 1, 1, 0) towards (1 − *E*_1_, *E*_2_, *ϵ*_3_, 1−*ϵ*_4_) and (*ϵ*_5_, 1 − *ϵ*_6_, 1 − *ϵ*_7_, *ϵ*_8_) where the *ϵ_i_* are below 0.5 but greater than zero Fig. 7d). Higher recombination rates (*ρ* ≥ 0.3) push *s*\* = 0.5 at *E*_2_ (Fig. 7b). Here, there is a predominance of (*ν_L_, ν_R_, ν_l_, ν_r_*) = (1,0.5,0,0.5) and (0,0.5,1,0.5) cells at equal frequencies (or (0.5, 1, 0.5, 0) and (0.5, 0, 0.5, 1) by symmetry). This arrangement is robust to recombination since the receptor locus is fixed at (*ν_R_, ν_r_*) = (0.5,0.5) and the ligand locus is either at (*ν_L_, ν_l_*) = (1,0) or (0,1) (or the ligand locus is (*ν_L_, ν_l_*) = (0.5,0.5) and the receptor is either at (*ν_R_, ν_r_*) = (1,0) or (0,1)). So pairing between these two cell types results in (1,0.5,0,0.5) and (0,0.5,1,0.5) offspring, whether recombination occurs or not. Note that this arrangement maintains some degree of asymmetry even with free recombination (*ρ* = 0.5). Even though both cell types produce both receptors, they produce the ligand asymmetrically (or vice versa). Cells on average are more likely to mate successfully between rather than within the two types of cells.

**Figure 7:**
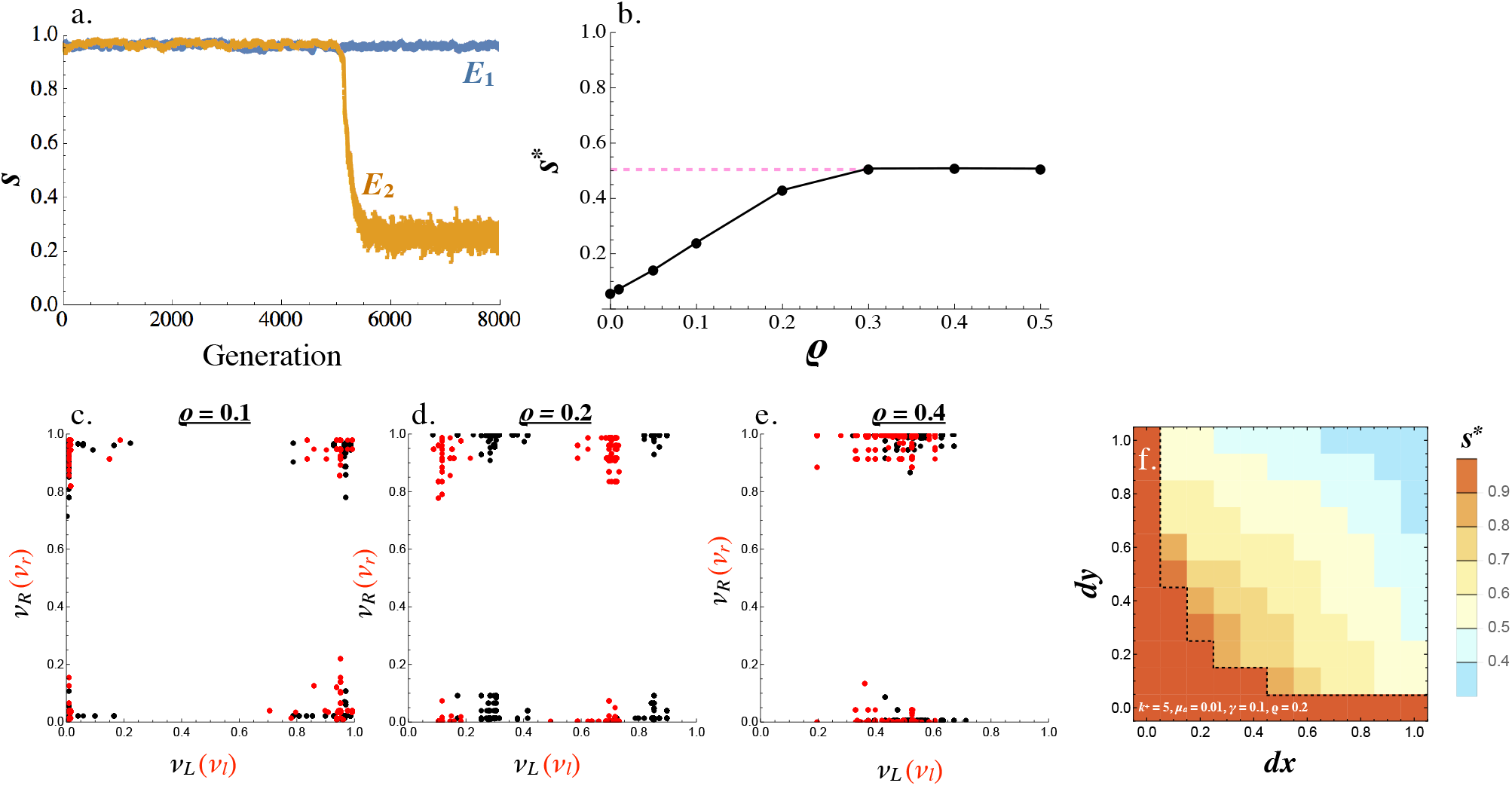
The effect of recombination on *E*_2_. (a) An example of evolution of the two signaling equilibria, *E*_1_ (for *k*^+^ =1) and *E*_2_ (for *k*^+^ = 5) given a fixed recombination rate *ρ* = 0.1. (b) Steady state *s*\* varies with the recombination rate. (c-d) Production rates of individual cells in the population for receptor-ligand pairs *L − R* (black) and *l − r* (red) for recombination rates (c) *ρ* = 0.1, (d) *ρ* = 0.2 and (e) *ρ* = 0.4. (f) Contour plot showing the steady state degree of symmetry (*s*\*) in a population with resident (*ν_R_, ν_L_, ν_r_, ν_l_*) = (1,1,0,0), given a recombination rate *ρ* = 0.2. Two mutations are introduced (1 − *dx*, 1, *dx*, 0) and (1,1 − *dy*, 0, *dy*) at rate *μ_a_* and their fate is followed until they reach a stable frequency. Other parameters used and simulation details are given in the Supplemental Material.

Similar to the case of no recombination, the invasion of *E*_1_ by *E*_2_ depends on the mutational process and parameter values. Fig. 7f shows the steady state symmetry measure in a population initially at (*ν_L_, ν_R_, ν_l_, ν_r_*) = (1, 1, 0, 0) when two mutations (1 − *dx*, 1, *dx*, 0) and (1,1 − *dy*, 0, *dy*) are introduced at low frequencies. Whether or not the mutants invade depends on the magnitude of the mutation in a similar way as in the case of no recombination (Fig. 5c versus Fig. 7f). However, the value of *s*\* now diverges from 0 reflecting the nonzero rate of recombination.

### 3.4 Evolution of linkage

In the analysis above, recombination between the ligand and receptor loci is fixed. However, the recombination rate itself can evolve. Let *ρ* undergo mutation at rate *μ_ρ_* so that the mutant *ρ*′ = *ρ*+*ε_ρ_* with *ε_p_* ~ *N*(0, *σ_ρ_*). In a diploid zygote, the rate of recombination is given by the average of the two recombination alleles, *ρ*_1_ and *ρ*_2_, carried by the mating cells. In this way, the recombination rate evolves together with the ligand and receptor production rates. We start with maximal recombination rate *ρ* = 0.5 and (*ν_L_, ν_R_, ν_l_, ν_r_*) = (1,1,0,0) for all cells and allow the recombination rate to evolve by drift for 1000 generation before we introduce mutation in the ligand and receptor loci.

The recombination rate evolves to *ρ*\* = 0 whenever *E*_2_ was reached from *E*_1_ in the non-recombination analysis. Under these conditions, tight linkage between receptor and ligand genes is favored (Fig. 8a). Furthermore, asymmetric signaling roles coevolve together with the recombination rate. The evolved trajectories of *s* and *ρ* depend on the strength of selection for asymmetric signaling. For example, when *k*^+^ is large (*k*^+^ = 10), signal asymmetry rapidly evolves; *s* moves away from 1 and this is followed by a sharp drop in the recombination rate (Fig. 8b). Eventually the population evolves asymmetric signaling roles (*s* in orange, Fig. 8b) and tight linkage (*ρ* in blue, Fig. 8b). These dynamics are similar when *k*^+^ is smaller (*k*^+^ = 3, Fig. 8c) and selection for asymmetry is weaker. However, it now takes longer for the asymmetric types to co-evolve (Fig. 8c). When selection for asymmetric signaling is even weaker (*k*^+^ =1, fig. 8d), no asymmetry evolves (*s* remains at 1) and the recombination rate fluctuates randomly between its minimum and maximum value as one would expect in the case of a neutral allele.

**Figure 8:**
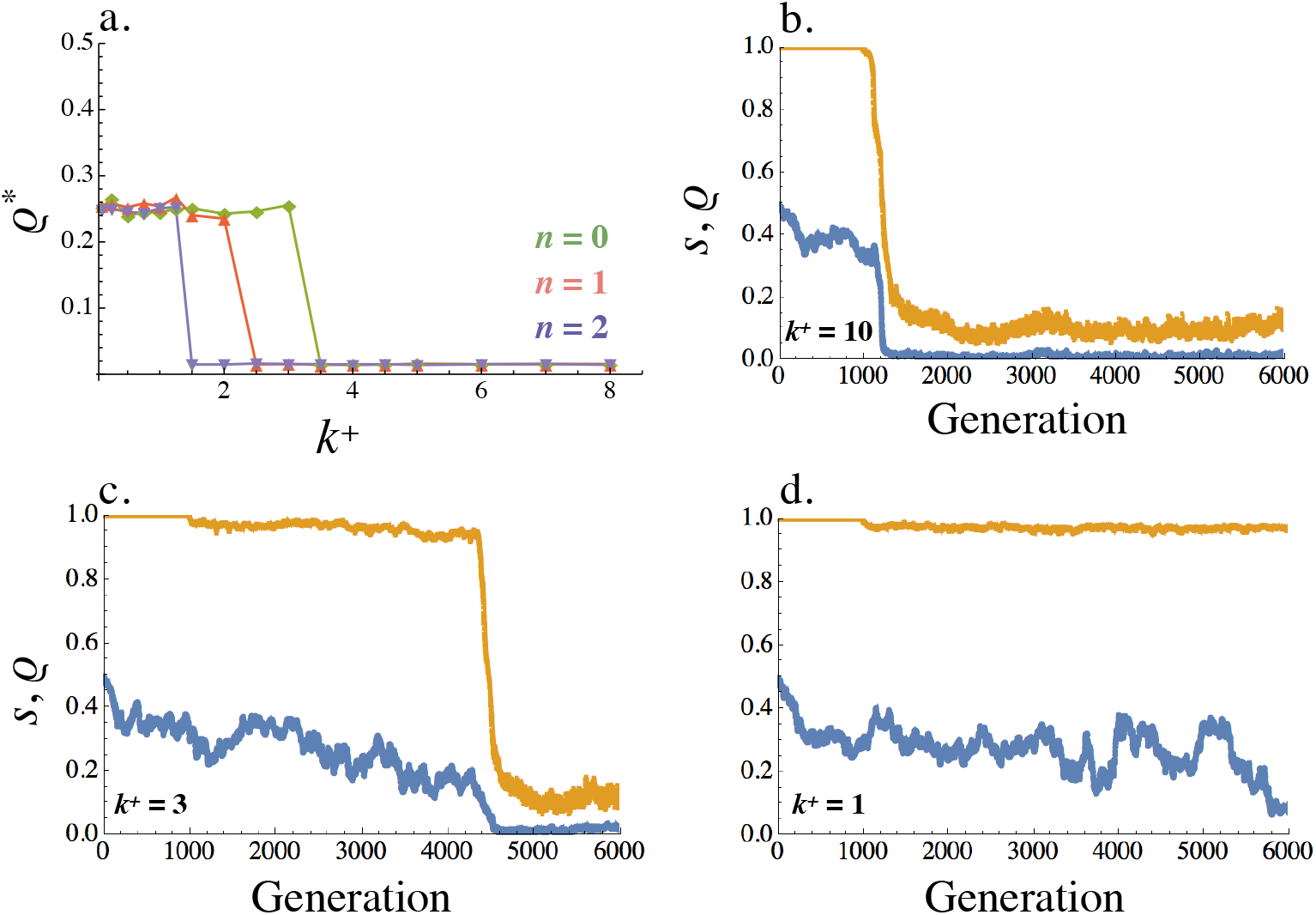
Equilibrium recombination rate *ρ*\*. (a) Averaged across the population, *ρ*\* varies with *k*^+^ (within cell binding rate) and *n* = 0,1,2 (cost of self-binding). (b-d) Evolution of the recombination rate *ρ* (blue) and signaling symmetry levels *s* (orange) for different within cell binding rates: (b) *k*^+^ = 10, (c) *k*^+^ = 3 and (d) *k*^+^ = 1. The recombination rate evolves under drift for the first 1000 generations, following which mutation at the ligand and receptor loci were introduced. When no asymmetry evolves the recombination rate fluctuates randomly between 0 and 0.5 (i.e. between its minimum and maximum value like a neutral allele). Other parameters used in simulations are given in the Supplemental Material.

## 4 Discussion

Explaining the evolution of mating types in isogamous organisms constitutes a major milestone in understanding the evolution of anisogamy and sexes [1, 3]. Mating type identity is determined by a number of genes that reside in regions of suppressed recombination and code for ligands and receptors that guide partner attraction and recognition, as well as genes that orchestrate cell fusion and postzygotic events [27, 8, 13, 12]. In this work we show that an asymmetry in ligand and receptor production evolves as a response to selection for robust gamete communication and swift mating. Furthermore, the same conditions favoring asymmetric signaling select for tight linkage between the receptor and ligand genes. Our findings indicate that selection for asymmetric signaling roles could have played an important role in the early evolution of gamete differentiation and identity.

We investigated the evolution of mating type roles by considering two types of ligand and receptor in individual cells. Gene duplication followed by mutation is a well established route to novelty evolution [38, 39, 40], and could explain the co-existence of two pairs of ligand and receptor in our system. Alternatively, individual cells could produce multiple ligands and receptors which evolve independently, as is the case in some basidiomycete fungi [41]. The production rate of the two types of ligand (and receptor) in our system is subject to mutation and selection so that the amount of expressed ligand (and receptor) of each kind is modulated quantitatively. In this way we were able to explicitly express the likelihood of mating as a function of the amount of free and bound molecules on the cell membrane and the ability of cells to accurately read their partner’s signal. This framework also allowed us to quantitatively follow the evolution of ligand and receptor production in mating cells for the first time.

We found that the ligand-receptor binding rate within a cell (*k*^+^) is key in the evolution of asymmetric signaling roles (Fig. 3, 4). *k*^+^ holds an important role because it dictates the rate at which free ligand and receptor molecules are removed from the cell surface. In addition, *k*^+^ determines the amount of intracellular signal that interferes with the ability of cells to interpret incoming signal. Although in theory cells could avoid self-binding (by reducing k^+^ to zero), there is likely to be a strong association of the within-cell and between-cell binding affinities. So reductions in *k*^+^ are likely to have knock-on costs in reducing *k_b_* as well. An extreme example is the case of locally diffusible signals (Fig. 1), such as those used by ciliates and yeasts to stimulate and coordinate fusion [29, 42]. Here binding affinities between and within cells are inevitably identical (since the ligand is not membrane bound). Work in yeast cells has shown that secreted ligands utilized for intercellular signaling during sex are poorly read by cells that both send and receive the same ligand [32]. In the case of strictly membrane bound molecules avoiding self-binding could also be an issue as it requires a ligand and receptor pair that bind poorly within a cell without compromising intercellular binding. For example, choosy budding yeast gametes (which are better at discriminating between species) take longer to mate [43]. It would be interesting to further study these trade-offs experimentally.

We never observed the co-existence of a symmetric “pansexual” type with asymmetric self-incompatible types. The two steady states consist of either a pansexual type or two mating types with asymmetric signaling roles. This follows from the requirement for strong selection to initiate evolution towards asymmetric signaling roles, and could explain why the co-existence of mating types with pansexuals is rare in natural populations [11, 12]. This is in contrast to previous models where pansexual types were very hard to eliminate due to negative frequency dependent selection [16, 44, 24]. For example, in the case of the mitochondrial inheritance model, uniparental inheritance raises mean population fitness, not only in individuals that carry genes for uniparental inheritance but also for pansexual individuals (benefits “leak” to biparental individuals)[24, 45].

A similar pattern is seen with inbreeding avoidance because the spread of self-incompatibility reduces the population mutation load, and so reduces the need for inbreeding avoidance [16]. These dynamics are reversed in the present model where there is positive frequency dependent selection. The spread of asymmetric signalers generates stronger selection for further asymmetry (Fig. 3, 4). This also occurs when there is recombination (Fig. 7, 8). Even though recombination between the two asymmetric types generates symmetric recombinant offspring, these are disfavored and eliminated by selection. These observations suggest that the mitochondrial inheritance and inbreeding avoidance models are unlikely to generate strong selection for suppressed recombination which is the hallmark of mating types. Finally, it would be interesting to explore how the reinstatement of recombination could be a route back to homothallism which is a state derived from species with mating types [12].

Mating type identity in unicellular eukaryotes is determined by mating type loci that typically carry a number of genes [27, 11]. Suppressed recombination at the mating type locus is a common feature across the evolutionary tree [8]. Our work predicts the co-evolution of mating type specific signaling roles and suppressed recombination with selection favoring linkage between loci responsible for signaling and an asymmetry in signaling roles. This finding suggests that selection for asymmetric signaling could be the very first step in the evolution of tight linkage between genes that control mating type identity. In yeasts, the only genes in the mating type locus code for the production of ligand and receptor molecules [29]. These then trigger a cascade of other signals downstream that also operate asymmetrically. Evidence across species suggests that mating type loci with suppressed recombination are precursors to sex chromosomes [46, 47]. In this way our work provides crucial insights about the origin of sex chromosomes.

The framework developed here could be used together with recent efforts to understand numerous features of mating type evolution. For example, opposite mating type gametes often utilize diffusible signals to attract partners [48, 49]. The inclusion of long range signals such as those used in sexual chemotaxis will provide further benefits for asymmetric signaling roles and mating types [26]. Furthermore the number of mating types varies greatly across species and is likely to depend on the frequency of sexual reproduction and mutation rates [50]. Signaling interactions between gametes could also play a role in determining the number of mating types and reducing their number to only two in many species [27]. It would be interesting to use the framework developed here to study the evolution of additional ligands and receptor and their role in reaching an optimal number of mating types. Other important features such as the mechanism of mating type determination [12, 51] and stochasticity in mating type identity [52, 53, 54] could also be understood in light of this work.

Taken together our findings suggest that selection for swift and robust signaling interactions between mating cells can lead to the evolution of self-incompatible mating types determined at non-recombinant mating type loci. We conclude that the fundamental selection for asymmetric signaling between mating cells could be the very first step in the evolution of sexual asymmetry, paving the way for the evolution of anisogamy, sex chromosomes and sexes.

## 5 Methods

### 5.1 General model

We model *N* cells so that each cell is individually characterized by a ligand locus 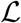 and a receptor locus 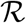. Two ligand genes at the locus 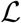 determine the production rates for two ligand types *l* and *L* given by *ν_l_* and *ν_L_*. Similarly, two receptor genes at the locus 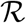 determine the production rates for the two receptor types *r* and *R* given by *ν_r_* and *ν_L_*. The two ligand and receptor genes in our model could could arise from duplication followed by mutation that leaves two closely linked genes that code for different molecules. In our computational set-up each cell is associated with production rates *ν_l_, ν_L_, ν_r_* and *ν_R_* where we assume a normalized upper bound so that *ν_l_* + *ν_L_* < 1 and *ν_r_* + *ν_R_* < 1.

The steady state concentrations for *L, R*, and *LR* are computed by setting 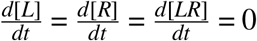 in Eq. (1–3) and solving the resulting quadratic equations. This leads two solutions only one of which gives positive concentrations. It follows that there is a unique physical solution to our system, which is what we use to define the probability of mating in our numerical simulations.

The program is initiated with *ν_L_* = *ν_R_* = 1 and *ν_l_* = *ν_r_* = 0 for all cells (unless otherwise stated, see Section 5.4). We introduce mutation so that the ligand and receptor production rates of individual cells mutate independently with probability *μ*. A mutation event at a production gene changes the production rate by an increment *ϵ* where *ϵ* ~ *N*(0, *σ*). Mutation events at the different genes *l, L, r* and *R* are independent of one another. If *ν_l_* + *ν_L_* > 1 or *ν_l_ + ν_L_* > 1 the production rates are renormalized so their sum is capped at 1. If a mutation leads to a production rate below 0 or above 1 it is ignored and the production rate does not change.

We implement mating by randomly sampling individual cells. The probability that two cells mate is determined by their ligand and receptor production rates as defined in Eq. (9) in the main text. We assume that *K* takes a large value relative to *W*_12_*W*_21_ so that *P* is linear in *W*_12_*W*_21_. Because the absolute value for *W*_12_*W*_21_ varies greatly between parameter sets, and what we are interested in is the relative change in *W*_12_*W*_21_ when signaling levels change, we chose *K* to be equal to the maximum value *W*_12_*W*_21_ can take for a given choice of *γ*, *k*^+^, *k*^−^ and *k_b_*. Sampled cells that do not mate are returned to the pool of unmated cells. This process is repeated until *M* = *N*/2 cells have successfully mated. This produces *N*/4 pairs of cells each of which gives rise to two offspring. These are sampled with replacement until the population returns to size *N*. We assume that a mutation-selection balance has been reached when the absolute change in s, defined in Eq. (10) in the main text, between time steps *t*_1_ and *t*_2_ is below *ϵ* = 10^−5^ across *t*_2_ − *t*_1_ = 100. Certain parameter sets resulted in noisy steady states and were terminated following 10^5^ generations. The numerical code keeps track of all production rates for individual cells over time.

### 5.2 Adaptive dynamics

We model adaptive dynamics by initiating the entire population at state (*ν_L_, ν_R_, ν_l_, ν_r_*)_*res*_ and introducing a mutant (*ν_L_, ν_R_, ν_l_, ν_r_*)_*mut*_ at low frequency *μ_a_*. We allow the population to evolve according to the life cycle introduced in the main text and record the frequency of the resident and mutant type when a steady state is reached. For the purposes of Fig. 5, the resident type is set to (*ν_L_, ν_R_, ν_l_, ν_r_*)_*res*_ and two mutants (*ν_L_, ν_R_, ν_l_, ν_r_*)_mut_1__ and (*ν_L_, ν_R_, ν_l_, ν_r_*)_*mut*_2__ are introduced both at frequency *μ_a_*. In this case we track the frequencies of the resident and both mutants until steady state is reached. We define steady state as the point where the average value of *s* in the population between time steps *t*_1_ and *t*_2_ is below *ϵ* = 10^−7^ across *t*_2_ − *t*_1_ = 100. The populatiomn always reached steady state.

### 5.3 Recombination

We implement recombination by considering a modifier 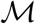 that lies between the ligand and receptor loci 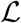 and 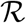. That is, we assume that the two ligand genes and two receptor genes are tightly linked on the ligand and receptor locus 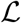 and 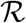 respectively, and only model recombination between the two loci. For simplicity, we assume that 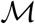 determines the physical distance between 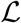 and 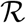 so that the distances 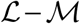 and 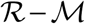 are the same. The modifier 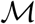 determines the rate of recombination between the ligand and receptor loci quantitatively by determining *ρ_M_*, the probability of a single recombination event following mating. Consider for example two individuals whose ligand and receptor production rates and recombination rate are determined by the triplets *R*_1_ − *M*_1_ − *L*_1_ and *R*_2_ − *M*_2_ − *L*_2_, the possible offspring resulting from such a mating are given by,

1. *R*_1_ − *M*_1_ − *L*_2_ and *R*_2_ − *M*_2_ − *L*_1_ with probability (1 − *ρM*_1,2_)^2^ – equivalent to no recombination event
2. *R*_1_ − *M*_2_ − *L*_1_ and *R*_2_ − *M*_1_ − *L*_2_ with probability 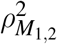 – equivalent to two recombination events
3. *R*_1_ − *M*_2_ − *L*_2_ and *R*_2_ − *M*_1_ − *L*_1_ with probability *ρM*_1,2_(1 − *ρM*_1,2_) – equivalent to one recombination event
4. *R*_1_ − *M*_1_ − *L*_2_ and *R*_2_ − *M*_2_ − *L*_1_ with probability *ρM*_1,2_(1 − *ρM*_1,2_) – equivalent to one recombination event

where 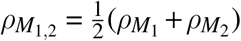 is the joint recombination rate when cell_1_ and cell_2_ with recombination rates *ρM*_1_ and *ρM*_2_ respectively mate.

We allow mutation at the recombination locus at rate *μ_ρ_* independently of the ligand and receptor loci. A mutation event leads to a new recombination rate so that 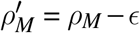 for *ϵ* ~ *N*(0, *σ_ρ_*). We assume that the mutation-selection balance has been reached when the absolute change in *s*, defined in Eq. (10) in the main text, and the change in the average recombination rate between time steps *t*_1_ and *t*_2_ is below *ϵ* = 10^−5^ across *t*_2_ − *t*_1_ = 100.

### 5.4 Methods and parameters used for simulated figures

**Fig. 4**

**(a)**: Individual simulations following the trajectory of *s* over time. Population is initiated at (*ν_L_, ν_R_, ν_l_, ν_r_*) = (1, 1, 0, 0) and *ρ* = 0 for all cells at time 0. *μ* = 0.01 for all ligand and receptor genes and *μ_r_* = 0. *σ* = 0.1, *γ* = 0.1, *k*^−^ =1, *n* =1, *k_b_* = *k*^+^/*k*^−^. *k*^+^ =1 for *E*_1_ trajectory and 5.0 for *E*_2_ trajectory. Population size *N* = 1000 and number of cells allowed to mate *M* = *N*/2.

**(b-c)**: Parameters as for (a) with *k*^+^ = 5.0. Each dot is represents an individual cell in the simulation.

**(d)**: Parameters used as for (a) with varying *k*^+^ and *γ* as indicated in the figure. Simulation was run until a steady state was reached and the value of *s*\* was averaged over the last 1000 time steps to account for noise.

**(e)**: Parameters used as for (a), varying *k_b_* and *n* as indicated in the figure. *k*^+^ was also varied here and the value of *k*^+^ beyond which *E*_2_ evolved at the expense of *E*_1_ was noted (the y-axis value).

**Fig. 5**

Adaptive dynamics simulations following the frequency of two mutants (*ν_L_, ν_R_, ν_l_, ν_r_*) = (1 − *dx*, 1, *dx*, 0) and (*ν_L_, ν_R_, ν_l_, ν_r_*) = (1,1 − *dy*, 0, *dy*) introduced at frequency *μ_a_* (indicated on figure) in a resident population with (*ν_L_, ν_R_, ν_l_, ν_r_*) = (1, 1, 0, 0). The frequency of the resident and two mutants at steady state was recorded and the heat maps show the average steady state value of *s*\* for 20 independent repeats. Parameters used: *γ* = 0.5, *k*^−^ =1, *n* =1, *k_b_* = *k*^+^/*k*^−^, *N* = 10000, *M* = *N*/2.

**Fig. 6**

**Joint evolution of receptor and ligand asymmetry.** Contour plots show the equilibrium frequency of a resident with production rates (*ν_L_, ν_R_, ν_l_, ν_r_*)_*res*_ = (1 − *dx*, 1, *dx*, 0) (a) (*ν_L_, ν_R_, ν_l_, ν_r_*)_*res*_ = (0.5 − *dx*, 0.5,0.5 + *dx*, 0.5) (b), following a mutation (*ν_L_, ν_R_, ν_l_, ν_r_*)_*mut*_ = (1,1 − *dy*, 0, *dy*) (a) and (*ν_L_, ν_R_, ν_l_, ν_r_*)_*mut*_ = (0.5,0.5 − *dy*, 0.5,0.5 + *dy*) (b). The mutant is introduced at a frequency *μ_a_* = 0.01. Other parameters used and simulations details are given in the Supplemental Material.

**Fig. 7**

**(a):** Individual simulations following the trajectory of *s* over time. Population is initiated at (*ν_L_, ν_R_, ν_l_, ν_r_*) = (1, 1, 0, 0) and *ρ* = 0.1 for all cells at time 0. *μ* = 0.01 for all ligand and receptor genes and *μ_r_* = 0. *σ* = 0.1, *γ* = 0.5, *k*^−^ = 1, *n* =1, *k_b_* = *k*^+^/*k*^−^. *k*^+^ = 1.0 for *E*_1_ trajectory and 5.0 for *E*_2_ trajectory. Population size *N* = 1000 and number of cells allowed to mate *M* = *N*/2.

**(b)**: Parameters as in (a) but varying *ρ* as indicated in the figure and using *k*^+^ = 3.0. The y axis shows the steady state value of *s* averaged over 1000 steps after steady state has been reached.

**(c-e)**: Parameters as for (a) with *k*^+^ = 5.0 and recombination rate *ρ* as shown in each figure. Each dot is represents an individual cell in the simulation.

**(f):** Parameters as for (a) with *k*^+^ = 5, *μ_b_* = 0.01, *ρ* =0.2 and *N*= 10000. The heat maps show the value of *s*\* at steady state averaged over 20 repeats. Heat map was obtained in the same way as Fig. 5.

**Fig. 8**

**(a):** Population is initiated at (*ν_L_, ν_R_, ν_l_, ν_r_*) = (1, 1, 0, 0) and *ρ* = 0.5 for all cells at time 0. *μ* = 0.01 for all ligand and receptor genes and *μ_p_* = 0.01. *σ* = *σ_ρ_* = 0.1, *γ* = 0.5, *k*^−^ =1, *k_b_* = *k*^+^/*k*^−^. *k*^+^ and *n* vary as shown in the plot. The y axis shows the steady state value of *ρ* averaged over 1000 steps after steady state has been reached. Population size *N* = 1000 and number of cells allowed to mate *M* = *N*/2.

**(b-d)**: Parameters as in (a) with *k*^+^ varied as shown in the individual plots.

## 6 Acknowledgements

This research was funded by an Engineering and Physical Sciences Research Council Fellowship (EP/L50488/) and HFSP Long Term Fellowship to ZH, and by grants from the Engineering and Physical Sciences Research Council (EP/F500351/1, EP/I017909/1, EP/K038656/1) and the Natural Environment Research Council (NE/R010579/1) to AP.

---

**Figure S1 Steady state concentrations in individual cells.** Steady state concentration of the ligand *L* and receptor *R* in individual cells when varying the ligand and receptor production rates *ν_L_* and *ν_R_* for *k*^+^/*k*^−^ = 10 (a) and *k*^+^/*k*^−^ = 0.1 (b). (c-d) show the concentration of ligand-receptor complexes for the same parameter variations. Other parameters used: *γ_R_* = *γ_L_* = *γ_LR_* = 0.1.

**Figure S2 The role of mutation rates.** The threshold value of *k*^+^, beyond which *E*_2_ becomes stable against *E*_1_, plotted versus *n* which dictates the cost of self-binding for *μ* = 0.1and *μ* = 0.001 to show that lower mutation rates require more stringent conditions for the evolution of signaling asymmetry. Population is initiated at (*ν_L_, ν_R_, ν_l_, ν_r_*) = (1, 1, 0, 0) and *ρ* = 0 for all cells at time 0. μ = 0.01 for all ligand and receptor genes and *μ_ρ_* = 0. *σ* = 0.1, *γ* = 0.5, *k*^−^ =1, *k_b_* = *k*^+^/*k*^−^. Population size *N* = 1000 and number of cells allowed to mate *M* = *N*/2.

**Figure S3 Synergy and competition between the production rates of the two ligands (and receptors).** Steady state signaling asymmetry *s*\* against the intracellular binding rate *k*^+^ for *ν_R_* + *ν_r_* < *α* and *ν_L_* + *ν_l_* < *α* for different values of a. *α* > 1 indicates synergy and *α* < 1 indicates competition between the two types of ligands (and receptors). For *α* = 0.75 the population only evolves asymmetric signaling for large values of *k*^+^(*k*^+^ = 5). In this case *s*\* is maximum at 0.75 since the sum of the two production rates cannot exceed 0.75. Population is initiated at (*ν_L_, ν_R_, ν_l_, ν_r_*) = (1, 1, 0, 0) and *ρ* = 0 for all cells at time 0. *μ* = 0.01 for all ligand and receptor genes and *μ_ρ_* = 0. *σ* = 0.1, *γ* = 0.5, *k*^−^ = 1, *k_b_* = *k*^+^/*k*^−^. Population size *N* = 1000 and number of cells allowed to mate *M* = *N*/2.

## References

[1] J. Lehtonen, H. Kokko, and G. A. Parker. What do isogamous organisms teach us about sex and the two sexes? Philosophical Transactions of the Royal Society B: Biological Sciences, 371(1706):20150532, 2016.

[2] G. A. Parker, R. R. Baker, and V. G. F. Smith. The origin and evolution of gamete dimorphism and the male-female phenomenon. Journal of Theoretical Biology, 36(3):529–553, 1972.

[3] J. R. Randerson and L. D. Hurst. The uncertain evolution of the sexes. Trends in Ecology and Evolution, 16(10):571–579, 2001.

[4] R. F. Hoekstra. The evolution of sexes. In S. C. Stearns, editor, The evolution of sex and its consequences, pages 59–91. Birkhauser Verlag, Basel, 1987.

[5] G. Bell. The evolution of anisogamy. Journal of Theoretical Biology, 73(2):247–270, 1978.

[6] B. Charlesworth. The population genetics of anisogamy. Journal of Theoretical Biology, 73(2):347–57, 1978.

[7] S. Branco, H. Badouin, R C. Rodríguez de la Vega, J. Gouzy, F. Carpentier, G. Aguileta, S. Siguenza, J. T. Brandenburg, M. A. Coelho, M. E. Hood, and T. Giraud. Evolutionary strata on young mating-type chromosomes despite the lack of sexual antagonism. Proceedings of the National Academy of Sciences, 114(27):7067–7072, 2017.

[8] S. Branco, F. Carpentier, R. C. R. De La Vega, H. Badouin, A. Snirc, S. Le Prieur, M. A. Coelho, D. M. De Vienne, F. E. Hartmann, D. Begerow, M. E. Hood, and T. Giraud. Multiple convergent supergene evolution events in mating-type chromosomes. Nature Communications, 9(1):2000, 2018.

[9] S. Ahmed, J. M. Cock, E. Pessia, R. Luthringer, A. Cormier, M. Robuchon, L. Sterck, A. F. Peters, S. M. Dittami, E. Corre, M. Valero, J. M. Aury, D. Roze, Y. Van De Peer, J. Bothwell, G. A. B. Marais, and S. M. Coelho. A haploid system of sex determination in the brown alga *ectocarpus sp*. Current Biology, 24(17):1945–1957, 2014.

[10] J. A. Fraser, S. Diezmann, R. L. Subaran, A. Allen, K. B. Lengeler, F. S. Dietrich, and J. Heitman. Convergent evolution of chromosomal sex-determining regions in the animal and fungal kingdoms. PLoS Biology, 2(12):e384, 2004.

[11] S. Billiard, M. López-Villavicencio, M. E. Hood, and T. Giraud. Sex, outcrossing and mating types: unsolved questions in fungi and beyond. Journal of Evolutionary Biology, 25(6):1020–38, 2012.

[12] S. Billiard, M. López-Villavicencio, B. Devier, M. E. Hood, C. Fairhead, and T. Giraud. Having sex, yes, but with whom? Inferences from fungi on the evolution of anisogamy and mating types. Biological Reviews of the Cambridge Philosophical Society, 86(2):421–42, 2011.

[13] N. Perrin. What uses are mating types? The “developmental switch” model. Evolution, 66(4):947–56, apr 2012.

[14] D. Charlesworth and B. Charlesworth. The evolution and breakdown of S-allele systems. Heredity, 43(1):41–55, 1979.

[15] M. K. Uyenoyama. On the evolution of genetic incompatibility systems. III. Introduction of weak gametophytic self-incompatibility under partial inbreeding. Theoretical Population Biology, 34(1):47–91, aug 1988.

[16] T. L. Czárán and R. F. Hoekstra. Evolution of sexual asymmetry. BMC Evolutionary Biology, 4(1):34, 2004.

[17] Z. Hadjivasiliou, A. Pomiankowski, R.M. Seymour, and N. Lane. Selection for mitonuclear co-adaptation could favour the evolution of two sexes. Proceedings of the Royal Society B: Biological Sciences, 279(1734), 2012.

[18] J. R. Christie, T. M. Schaerf, and M. Beekman. Selection against heteroplasmy explains the evolution of uniparental inheritance of mitochondria. PLoS Genetics, 11(4):e1005112, 2015.

[19] L. D. Hurst and W. D. Hamilton. Cytoplasmic fusion and the nature of sexes. Proceedings of the Royal Society B: Biological Sciences, 247(1320):189–194, 1992.

[20] L. D. Hurst. Why are there only two sexes? Proceedings of the Royal Society B: Biological Sciences, 263:415–422, 1996.

[21] V. Hutson and R. Law. Four steps to two sexes. Proceedings of the Royal Society B: Biological Sciences, 253(1336):43–51, 1993.

[22] J. R. Christie and M. Beekman. Uniparental inheritance promotes adaptive evolution in cytoplasmic genomes. Molecular Biology and Evolution, 34(3):677–691, 2017.

[23] I. M. Hastings. Population genetic aspects of deleterious cytoplasmic genomes and their effect on the evolution of sexual reproduction. Genetics Research, 59(3):215–25, 1992.

[24] Z. Hadjivasiliou, N. Lane, R. M. Seymour, and A. Pomiankowski. Dynamics of mitochondrial inheritance in the evolution of binary mating type and two sexes. Proceedings of the Royal Society B: Biological Sciences, 280(1769):20131920, 2013.

[25] A. J. Wilson and J. Xu. Mitochondrial inheritance: Diverse patterns and mechanisms with an emphasis on fungi. Mycology, 3(2):158–166, 2012.

[26] Z. Hadjivasiliou, Y. Iwasa, and A. Pomiankowski. Cell - Cell signalling in sexual chemotaxis: A basis for gametic differentiation, mating types and sexes. Journal of the Royal Society Interface, 12(109), 2015.

[27] Z. Hadjivasiliou and A. Pomiankowski. Gamete signalling underlies the evolution of mating types and their number. Philosophical Transactions of the Royal Society B: Biological Sciences, 371(1706), 2016.

[28] H. W. Kuhlmann, C. Brünen-Nieweler, and K. Heckmann. Pheromones of the ciliate *Euplotes octocarinatus* not only induce conjugation but also function as chemoattractants. The Journal of Experimental Zoology, 277(1):38–48, 1997.

[29] L. Merlini, O. Dudin, and S. G. Martin. Mate and fuse: how yeast cells do it. Open Biology, 3(3):130008, 2013.

[30] I. Maier. Gamete orientation and induction of gametogenesis by pheromones in algae and plants. Plant, Cell & Environment, 16:891–907, 1993.

[31] Y. Tsubo. Chemotaxis and sexual behavior in *Chlamydomonas*. The Journal of Protozoology, 8(2):114–121, 1961.

[32] H. Youk and W. A. Lim. Secreting and sensing the same molecule allows cells to achieve versatile social behaviors. Science, 343(6171):1242782, 2014.

[33] C. Cappellaro, K. Hauser, V. Mrsa, M. Watzele, G. Watzele, C. Gruber, and W. Tanner. *Saccharomyces cerevisiae* a-and a-agglutinin: characterization of their molecular interaction. The EMBO Journal, 10(13):4081–4088, 1991.

[34] N. F. Wilson, J. S. O’Connell, M. Lu, and W. J. Snell. Flagellar adhesion between mt+ and mt-*Chlamydomonas* gametes regulates phosphorylation of the mt+-specific homeodomain protein GSP1. Journal of Biological Chemistry, 274(48):34383–34388, 1999.

[35] S. S. Phadke and R. A. Zufall. Rapid diversification of mating systems in ciliates. Biological Journal of the Linnean Society, 98(1):187–197, 2009.

[36] Z. Hadjivasiliou, G. L. Hunter, and B. Baum. A new mechanism for spatial pattern formation via lateral and protrusionmediated lateral signalling. Journal of the Royal Society Interface, 13(124), 2016.

[37] L. LeBon, T. V. Lee, D. Sprinzak, H. Jafar-Nejad, and M. B. Elowitz. Fringe proteins modulate Notch-ligand cis and trans interactions to specify signaling states. eLife, 3:e02950, 2014.

[38] O. Susumu. Evolution by Gene Duplication. Springer Science, New York, 1970.

[39] J. Zhang. Evolution by gene duplication: An update. Trends in Ecology and Evolution, 18(6):292–298, 2003.

[40] S. Magadum, U. Banerjee, P. Murugan, D. Gangapur, and R. Ravikesavan. Gene duplication as a major force in evolution. Journal of Genetics, 92(1):155–161, 2013.

[41] T. J. Fowler and L. J. Vaillancourt. Pheromones and pheromone receptors in *Schizophyllum commune* mate recognition: a retrospective of a half-century of progress, and a look ahead. In J. Heitman, J. T. Kronstad, and L. A. Casselton, editors, Sex in fungi: Molecular determination and evolutionary iImplications, pages 301–315. American Society of Microbiology (ASM) Press, Washington D.C., 2007.

[42] M. Sugiura, H. Shiotani, T. Suzaki, and T. Harumoto. Behavioural changes induced by the conjugation-inducing pheromones, gamone 1 and 2, in the ciliate *Blepharisma japonicum*. European Journal of Protistology, 46(2):143–9, 2010.

[43] D. W. Rogers, J. A. Denton, E. McConnell, and D. Greig. Experimental evolution of species recognition. Current Biology, 25(13):1753–1758, 2015.

[44] R. F. Hoekstra. On the asymmetry of sex: evolution of mating types in isogamous populations. Journal of Theoretical Biology, 98(3):427–451, 1982.

[45] J. R. Christie and M. Beekman. Selective sweeps of mitochondrial DNA can drive the evolution of uniparental inheritance. Evolution, 71(8):2090–2099, 2017.

[46] S. Geng, P. De Hoff, and J. G. Umen. Evolution of sexes from an ancestral mating-type specification pathway. PLoS Biology, 12:1–16, 2014.

[47] A. Menkis, D. J. Jacobson, T. Gustafsson, and H. Johannesson. The mating-type chromosome in the filamentous ascomycete *Neurospora tetrasperma* represents a model for early evolution of sex chromosomes. PLoS Genetics, 4(3):e1000030., 2008.

[48] Y. Tsuchikane, T. Fujii, M. Ito, and H. Sekimoto. A sex pheromone, protoplast release-inducing protein (PR-IP) inducer, induces sexual cell division and production of PR-IP in closterium. Plant and Cell Physiology, 46(9):1472–1476, 2005.

[49] P. Luporini, A. Vallesi, C. Miceli, and R. A. Bradshaw. Chemical signaling in ciliates. The Journal of Eukaryotic Microbiology, 42(3):208–12, 1995.

[50] G. W. A. Constable and H. Kokko. The rate of facultative sex governs the number of expected mating types in isogamous species. Nature Ecology and Evolution, 2(7):1168–1175, 2018.

[51] S. Vuilleumier, N. Alcala, and H. Niculita-Hirzel. Transitions from reproductive systems governed by two self-incompatible loci to one in fungi. Evolution, 67(2):501–516, 2013.

[52] Z. Hadjivasiliou, A. Pomiankowski, and B. Kuijper. The evolution of mating type switching. Evolution, 70(7):1569–1581, 2016.

[53] B. P. S. Nieuwenhuis and S. Immler. The evolution of mating-type switching for reproductive assurance. BioEssays, 38(11):1141–1149, 2016.

[54] B. P. S. Nieuwenhuis, S. Tusso, P. Bjerling, J. Stangberg, J. B. W. Wolf, and S. Immler. Repeated evolution of self-compatibility for reproductive assurance. Nature Communications, 9(1):1639, 2018.

